# DNA extraction and sequencing of the Mawangdui ancient cadaver protected by formalin

**DOI:** 10.1101/2020.05.06.081711

**Authors:** Hua-Lin Huang, Shikui Yin, Huifang Zhao, Chao Tian, Jufang Huang, Sihao Deng, Zhiyuan Li

**Author notes:** Correspondence should be addressed to Dr. Zhiyuan Li, Guangzhou Institutes of Biomedicine and Health, Chinese Academy of Sciences, 190 Kai Yuan Avenue, Science Park, Guangzhou 510530, China.

## Abstract

Mawangdui ancient Cadaver is the first wet corpse found in the world, which is famous for being immortal for over two thousands of years. After being unearthed, the female corpse was immersed in the formalin protective solution for more than 40 years. We used magnetic bead method and formalin fixed paraffing (FFPE) method to extract the DNA of the female corpse, respectively. PCR amplification, sanger sequencing, library building, high throughput sequencing (testing) and data processing were carried out on the DNA samples, and about 0.5% of the whole genome coverage sequencing data was obtained. Comparing the results of DNA trough two extraction and sequencing methods. We found that the FFPE and high throughput sequencing methods is better than others for DNA extraction of the ancient samples which were preserved in formalin, providing a guidance for dealing with formalin preserved ancient samples in the future.

## Introduction

The first wet corpse in the world was excavated in 1972 from the Mawangdui Han tomb, which is located in Changsha, Hunan Province, China (Fig. 1A)(*1*). When the female corpse of Tomb No. 1 was unearthed, her body was preserved in a mix fluid inside the four coffins. Her body was intact and completely covered with moistened skin, the muscles still allowed some joint movement, some blood vessels were clearly visible, the soft connective tissues were elastic, and the toeprints and fingerprints were clear (Fig. 1C)(*1*). This kind of corpse was unprecedented in the world (*2*). Therefore, it was defined as wet corpse, a new type of corpse, which is distinct from not only tanned and wax-preserved corpses but also dry corpses, such as mummy (*1, 3, 4*).

**Figure 1.**
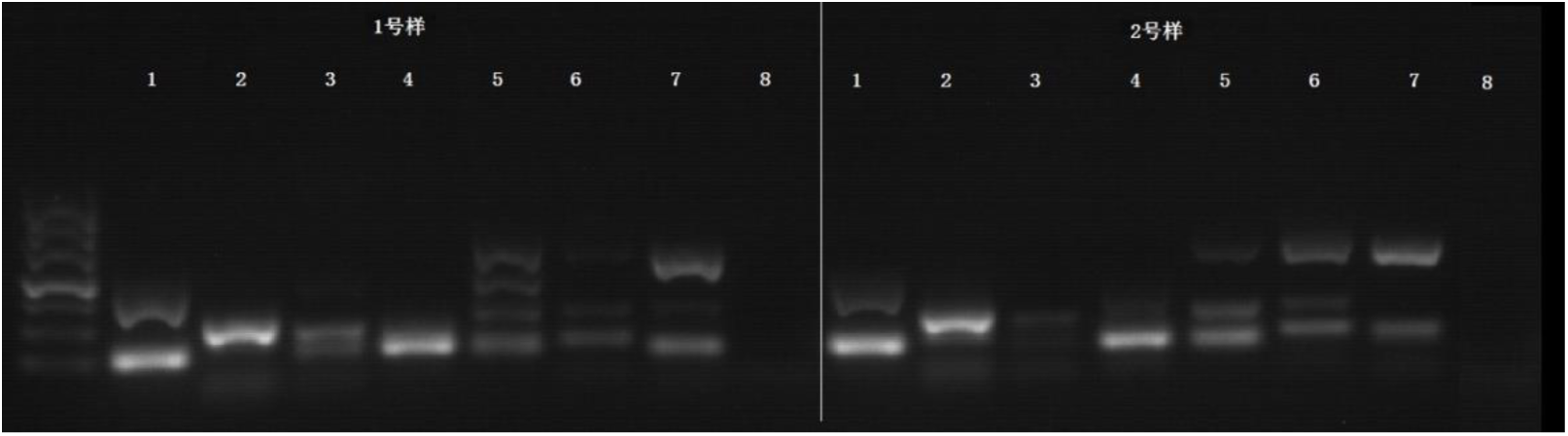
Mawangdui No.1 sample extracted by QIAamp-DNA-FFPE tissue method (left) and magnetic bead method (right)

Following with the unearthed Mawangdui wet corpse, more wet corpses were excavated. For examples, the Han Dynasty male corpse was unearthed in Jingzhou, Hubei Province in 1975, the Qing Dynasty female corpse was unearthed in Dangshan, Anhui Province in 2001, the Han Dynasty female corpse unearthed was in Lianyungang in 2002, and the Ming Dynasty female corpse was unearthed in Taizhou, Jiangsu Province in 2011.

After excavation, the Mawangdui wet corpse was stored in formalin-based fixative, of which the plexiglass coffin is held in the Hunan Provincial Museum. The preservation methods of other wet corpses are all based on formalin preservation solution. This is the first time to study the DNA extraction and sequencing of the wet corpse fixed by formalin, which is of great scientific value for other wet corpse research and social significance.

## Materials and methods

### Experimental materials and environment

The vertebrae of the female corpse of Mawangdui No.1 tomb (about 2cm long, 1.4cm wide, 1.4cm thick and 30g weight) were used in this experiment.

According to the requirement of ancient DNA research, in the ten thousand level positive pressure laminar flow chamber, hydrogen peroxide was used to clean the positive pressure super clean table, all the equipment, instruments and consumables, then did again by alcohol,. After removing the possible pollution, they were put into the table of the positive pressure super clean table. Before each experiment, the laminar flow chamber and the super clean table shall be subject to 0.5h ultraviolet irradiation before entering. Before laminar flow room, surgical hand washing operation shall be carried out. After wearing disposable gloves mask, hat and conjoined operation clothes in positive pressure dressing room. Finally, a pair of sterile rubber gloves shall be put on, and the operation room shall be entered 3 minutes later after wind drenching room. Only samples from the same source are taken each time. After each operation, alcohol, hydrogen peroxide and ultraviolet disinfection were carried out and the platform was sealed.

### DNA extraction

After we used a variety of DNA extraction methods in our preliminary experiment, we finally referred to the magnetic bead method and formalin fixed specimen embedding (FFPE) two extraction methods. The material is taken from the vertebrae of No.1 female corpse in Mawangdui. The bone tissue material used in each experiment was about 2G.

#### DNA extraction by improved DNA FFPE tissue method

The operation steps are as follows:

1. Add 25 ml ethanol (96% – 100%, analytical purity) to buffer aw1 in QIAamp DNA FFPE tissue, and 30 ml ethanol (96% – 100%) to buffer AW2. Mix the solution before each use, and it can be stored for one year at room temperature. If buffer al precipitates, it can be bathed in a 70 °C water bath pot, and can be used after it is melted.
2. Take the sample out of the refrigerator, put it at room temperature, use the scalpel to remove the 1-3mm tissue layer on the surface of the larger bone specimen, and then use the scalpel to cut up the required amount of tissue samples for use, the amount of each bone tissue sample is about 1cm^3^.
3. The primary cut bone tissue was put into a 15ml EP tube containing 8-10ml of 0.5m EDTA buffer. 37 ° C constant temperature oscillation, change EDTA every 12h, repeat 3 times.
4. Centrifugation at 13000 rpm at room temperature (15 – 25 ° C) for 2 minutes, using a pipette to remove the supernatant, taking care not to remove any precipitated particles, wash with deionized water twice, while transferring the precipitated particles to a 1.5 ml centrifuge tube.
5. Add 180 μ l ATL and 20 μ l proteinase K into the centrifuge tube, mix them with vortex, incubate them overnight at 56 ° C, dissolve the samples basically, and incubate them at 90 ° C for 1 hour after the samples fall to room temperature.
6. Cool the sample to room temperature, add 2 μ L RNaseA ((100 mg / ml), incubate for 2 minutes, centrifuge 1.5 ml centrifuge tube instantaneously, and centrifuge the liquid on the tube cover.
7. Add 200 μ l al buffer night and 200 μ l ethanol (99%) for premixing, mix the solution with a pipette, and then add it to the sample to be treated together, immediately and thoroughly vortex mix. Then add 200 μ l ethanol (99%), and immediately vortex until thoroughly mixed. After adding al buffer and ethanol, white precipitate will be produced. Centrifugate 1.5 ml centrifuge tube instantaneously and the liquid on the tube cover.
8. Carefully transfer all the cracking solution to the qiaamp extraction column of the 2 ml collecting tube, drop the solution to the center of the extraction column as far as possible, and centrifugate at 8000 rpm for 1 minute after closing the tube cover. Take out the centrifuge tube and put it into a new 2 ml collecting tube. The original collecting tube and the liquid in it can be discarded. Centrifuge again at 8000 rpm until there is no residual solution in the QIAamp extraction column.
9. Open the cover of the QIAamp extraction column and add 500 μ l of aw1 buffer to the center of the column. Cap the centrifuge tube and centrifuge at 8000 rpm for 1 minute. Take out the centrifuge tube and put it into a clean 2 ml collecting tube. The original collecting tube can be discarded.
10. Carefully open the cover of the QIAamp extraction column and add 500 μ l of AW2 buffer to the core of the extraction column. After tightening the centrifuge tube cover, centrifuge at 8000 rpm for 1 minute. Take out the centrifuge tube and put it into a new 2 ml collecting tube. The original collecting tube and the liquid in it can be discarded. Avoid contact with the QIAamp extraction column.
11. Carefully place the qiaamp extraction column in a new 2 ml collecting tube, centrifuge at 14000 rpm for 1 minute, and try to dry the column membrane to avoid the interference of residual ethanol in the effluent with subsequent nucleic acid amplification experiments.
12. Carefully place the qiaamp extraction column in a new 1.5 ml centrifuge tube, discard the original collection tube and the liquid in it. Carefully open the cover of qiaamp extraction column and add about 50 μ l of buffer ate which has been balanced at room temperature to the column center.
13. Cover the extraction column, leave it at room temperature for 5 minutes, centrifugation at 14000 rpm for 1 minute, and discard the qiaamp extraction column. Sub Pack 2 ul for concentration determination. DNA samples were stored in – 80 °C refrigerator for a long time.

#### Extraction of tissue genomic DNA by improved magnetic bead method

The extraction steps are as follows:

1. Remove the sample from the refrigerator and place it at room temperature. Use a scalpel to remove the 1-3mm tissue layer on the upper surface of the larger bone specimen, and then use the surgical scissors to cut the required amount of tissue samples, and cut them into pieces. Each time, the amount of bone tissue samples is about 1cm^3^.
2. The primary cut bone tissue was put into a 15ml centrifuge tube containing 8-10ml of 0.5m EDTA buffer. 37 ° C constant temperature oscillation, change EDTA every 12 hours, repeat 3 times. At room temperature (15 – 25 ° C)
3. Centrifugation at 13000 rpm for 2 minutes, use a pipette to remove the supernatant, handle with care, do not remove any particle precipitation, wash twice with deionized water, and transfer the precipitated particles to a 1.5 ml centrifuge tube.
4. Add 200 μ l of GHA tissue digestive solution, 20ul of DTT and 20 μ l of protease K, digest overnight at 56 °C, centrifuge at 13000 rpm for 1 minute, take the supernatant with a pipette and put it into a new 1.5 ml centrifuge tube to remove the residual impurities. Add 4 μ l RNase A and leave at room temperature for 10 minutes.
5. Add 300 μ l of GHB cracking liquid, shake and mix well. Put 1.5 ml centrifuge tube at 75 °C, incubate for 15 minutes, during which it needs to be reversed and mixed 3 times, 5 times each time. Leave at room temperature for 5 minutes.
6. Add 350 μ l isopropanol, shake and mix for 10 seconds.
7. Add 30 μ l suspension g of magnetic beads, shake and mix for 1 minute, leave for 9 minutes in total, shake and mix for 1 minute every 3 minutes.
8. Place the centrifuge tube on the magnetic frame and let it stand for 30 seconds. After the magnetic bead is completely absorbed, use the pipette to carefully absorb the liquid.
9. Add 700 μ l of GDA buffer, shake and mix for 30 seconds.
10. Place the centrifuge tube on the magnetic frame and let it stand for 30 seconds. After the magnetic bead is completely absorbed, carefully absorb the liquid. Repeat these two steps once.
11. Add 700 μ l PWD rinse solution, shake and mix for 30 seconds.
12. Place the centrifuge tube on the magnetic frame and let it stand for 30 seconds. After the magnetic beads are completely absorbed, use the pipette to carefully absorb the liquid. Repeat these two steps once.
13. Put the centrifuge tube on the magnetic frame and dry at room temperature for 15 minutes to ensure that the ethanol is completely volatilized and clean, so as to avoid the influence of ethanol residue on the subsequent enzyme reaction.
14. Remove the centrifuge tube from the magnetic frame, add 50 μ l of eluent buffer TB, shake and mix, and incubate in a 56 °C water bath for 10 minutes, during which, mix upside down 3 times, 5 times each time.
15. Put the centrifuge tube on the magnetic frame again and let it stand for 2 minutes. After the magnetic beads are completely absorbed, carefully transfer the DNA solution to a new centrifuge tube and repack it with 2 ul for concentration determination. DNA samples were stored in −80 °C refrigerator for a long time.

### PCR amplification and Sanger sequencing

The OD value of the sample (a260 / 280) was measured by nanodrop 2000 spectro-photometer. PCR amplification and Sanger sequencing of the extracted ancient DNA were carried out. According to the literature reports of the identification of Czar’s remains, the ancient DNA of Jiaohe and Neanderthal, the corresponding identification primers were designed, which were four pairs of mitochondrial DNA and four pairs of nuclear DNA. Through NCBI genome database (http://www.ncbi.nlm.nih.gov/gene) and prime soft 5.0 (https://enterthegungeon.gamepedia.com/prime_prime), the target fragment of the primer was obtained, and the length and annealing temperature of the target fragment were deduced (Table 1).

**Table 1.**
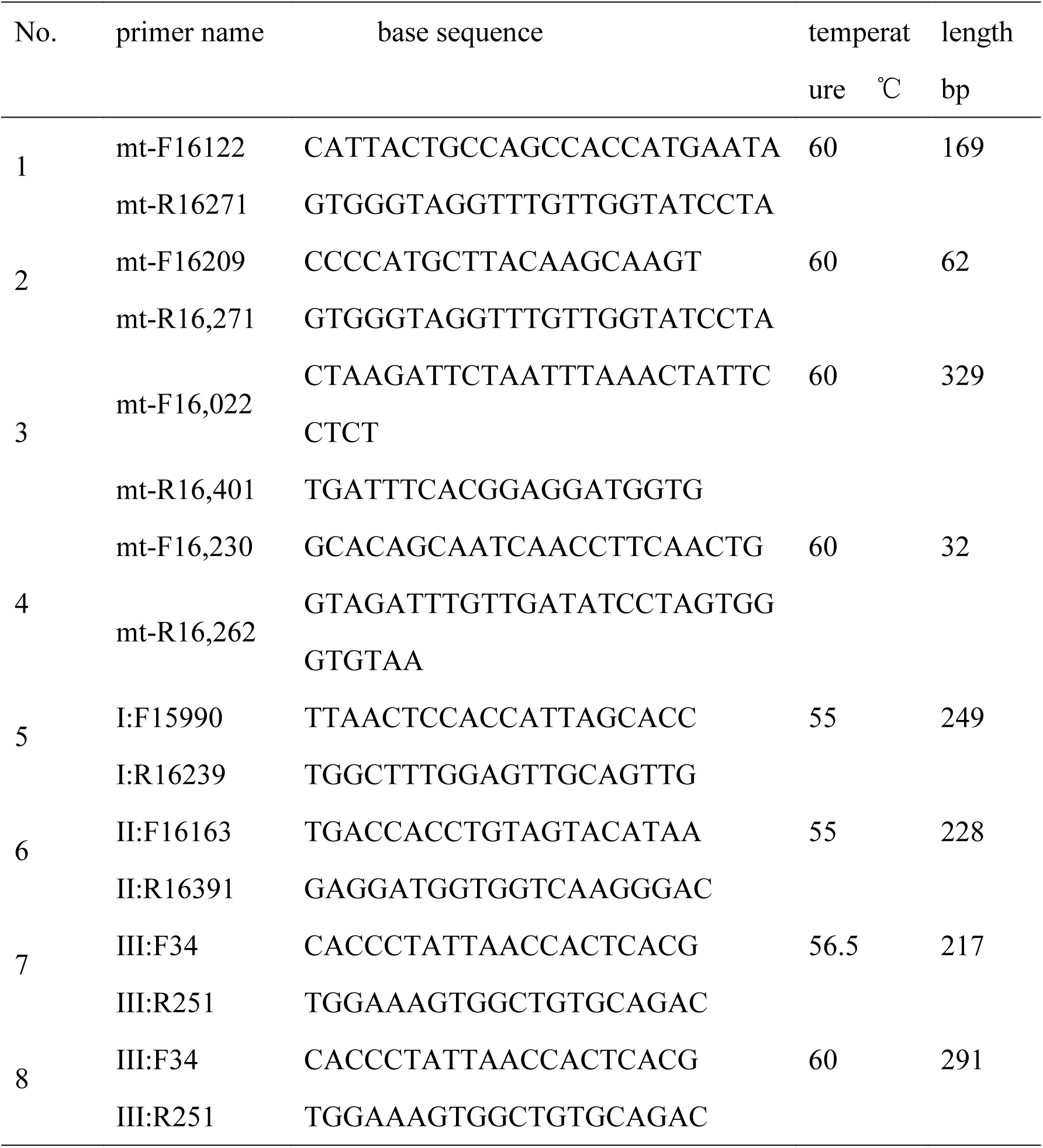
primers, target fragment length and annealing temperature

2% agarose gel electrophoresis detection with 120V constant pressure electrophoresis for 30 minutes; agarose gel into gel imaging system, open ultraviolet light, observe the results of running gum and take pictures, then clean surgical blades to cut the appropriate size strip under ultraviolet lamp indication to purify and recycle the product.

Sequencing analysis: the purified PCR products and primers were sequenced in the first generation. The sequencing primers were the above 8 PCR primers. The sequencing results were analyzed by chromas and DNAStar software, and the target gene standard sequence in NCBI database was sequenced.

### Library establishment and high throughput sequencing

After qubit was used to detect the concentration of DNA sample, DNA repair was carried out directly, without breaking and gel recovery, and 5’ phosphorylated flat ended DNA was produced, so that it could be connected with the specific aptamer of sequencing platform. The efficiency of library construction directly depends on the accuracy and efficiency of DNA end repair. The end repair mixture transforms the 5’ or 3’ sticky end into 5’ phosphorylated flat end DNA. In most cases, terminal repair is accomplished by the 5’ - 3’ polymerase activity of T4 DNA polymerase and the 3’ - 5’ exonuclease activity. T4 polynucleotide kinase ensures phosphorylation of the 5’ end of the flat ended DNA fragment for subsequent aptamer connection. A single prominent adenylate a needs to be added to the 3’ end of the fragment to pair with the prominent thymidylate t on the platform specific aptamer. In general, this step is catalyzed by a Klenow fragment with the lowest exonuclease activity from 3’ to 5’. T4 DNA ligase connects the double stranded aptamer with the repaired end of the library fragment, and then selectively removes the unconnected aptamer and aptamer dimer in the library according to the size of DNA. Size screening methods include agarose gel electrophoresis and separation.

Bridge PCR: adjust the above DNA sample to a suitable concentration and then add it to flowcell, so that one end of the sequence is combined with the existing short sequence on flowcell by complementary pairing, and then the bridge PCR can be officially started after the connection. The first amplification was carried out, and the sequence was added into double chain. Add NaOH strong alkaline solution to destroy the double strand of DNA, and then elute. Add buffer to make the reaction environment neutral, and then the free end of the sequence will be matched with the adjacent adpater. In the next round of PCR, in the process of PCR, the sequence will be bent into a bridge, and one round of bridge PCR can double the sequence amplification. In this way, we will get a cluster with the same sequence, which is generally called cluster.

Sequencing: using illunima3000 sequencing platform for high-throughput sequencing, when constructing the DNA library to be tested, using the paired end method to add all the binding sites of sequencing primers on the joints at both ends, and then adding special treatment (adding azide group and fluorescence group to the base part of DNA 3 to excite different colors) a, t, C and G bases. In the process of sequencing, only one base is added to the current sequencing chain. At this time, the sequencer will emit excitation light and scan fluorescence at the same time. Because all sequences in a cluster are the same, in theory, the fluorescence color emitted in the cluster should be the same. On this basis, add reagents to change the – N2 at the position of deoxyribose 3 to – Oh, and cut off some fluorescence groups, so that they will no longer emit fluorescence in the next round of reaction. In this way, the contents of the sequence can be measured in cycles. After the completion of the first round of sequencing, the template chain of the first round of sequencing is removed. In order to guide the regeneration and amplification of complementary chain in the original position, the paired end module is used to ensure that the synthesis of the second round of complementary chain reaches the template quantity for sequencing.

### Quality control of sequencing data

General quality control of sequencing data is shown in Figure 2, 2: after base recognition, the original image file (BCL) obtained by sequencing will be transformed into raw data in fastq format, and then the quality analysis of the original sequencing data will be carried out to evaluate whether the sequencing data is suitable for the next step of bioinformatics analysis, in which the quality analysis mainly includes sequencing quality and base composition analysis. Based on the analysis of the base composition and quality value of the data, the data are filtered according to the analysis results of raw data to remove the joint sequence and the polluted part. To remove too many low-quality base sequences, the standards of data filtering mainly include the following: 1. To remove the pollution of the connector, use adapterremoval (version 2.1.7) to remove the pollution of the 3’ end connector; 2. To filter the quality, use the sliding window method to filter the quality of the next step. The window size is set to 5 BP, and the step size is generally set to 1 BP, move one base forward each time, take 5 bases to calculate the average Q value of the window, if the Q value of the last base is ≤ 2, only the base before the position is reserved; if the average Q value of the window is ≤ 20, only the last base of the window and the previous base (6) are reserved. Q value is the abbreviation of base quality value, which is an important standard to evaluate the error rate of sequencing results. Q value is the rounding mapping result reflecting the base reading error rate P during sequencing, as shown in table 2.8. The data filtered by two terminal sequencing needs to be further screened, and the paired sequences need to be preserved to get clean data. After data filtering, make quality statistics on the obtained clean data, count the base content, alkali and base mass distribution, confirm the filtered data, and observe whether it meets the analysis requirements. If the analysis requirements are met, it is used for the next analysis. If the analysis requirements are not met, the data needs to be filtered again.

**Figure 2.**
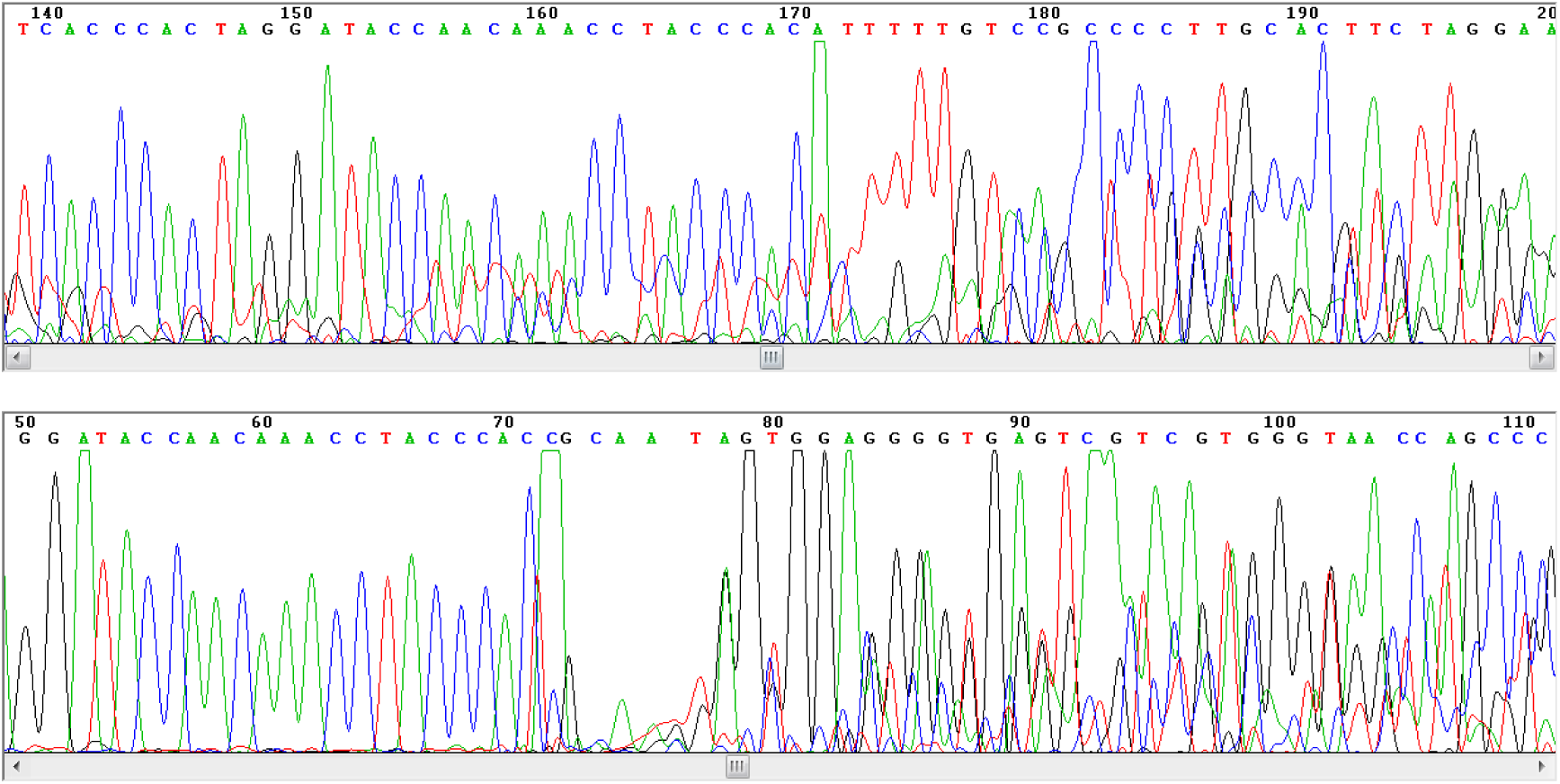
Partial sequencing peaks of sample 1 and sample 2

Quality control of ancient DNA sequencing data: damage detection shall be carried out on the acquired data, mainly to detect the change of cytosine (c) into thymine (T), guanine (g) into adenine (a), CPG into TPG and CpG into CPA, so as to select sequencing data conforming to the characteristics of ancient DNA sequence for further analysis (7).

### SNP analysis

Single nucleotide polymorphism (SNP) mainly refers to the DNA sequence polymorphism caused by the variation of single nucleotide at the genomic level, including the transformation and transversion of single base (11). Using gatk software to detect SNP, using haplotypecaller program (stand exit conf set to 10, stand call conf set to 30) to obtain the mutation sites of the samples, extract the SNP mutation sites, in order to ensure the reliability of the sites, but also to take into account the factors that are difficult to sequence ancient DNA, filter the SNP sites, the filtering standards are as follows: (1) Depth(DP) ≥ 4; (2) Fisher test of strand bias (FS) ≤ 60; (3) HaplotypeScore ≤ 13.0; (4) Quality Depth(QD) ≥ 2; (5) Mapping Quality (MQ) ≥ 40; (6) ReadPosRankSum ≥ −8.0; (7) MQRankSum ≥ −12.5。

In order to further study the characteristics of mutations, we can see the preference of different types of SNP bases before and after mutations according to the types of bases before and after mutations. The SNP results obtained from the analysis were statistically analyzed.

The SNP loci were annotated by anovar software (12). The annotation software integrates multiple databases and analysis resources, including refgene (13), cycloband, exac (exome aggregation consortium), nhlbi-esp6500 (NHLBI go exome sequencing project), 1000 genes project, Cosmo (catalog of social musings in cancer), clinvar, OMIM (Online Mendelian inheritance in) Man (14), and SIFT (sorting assistant from assistant) (15), polyphen2 (16), mutation aster, mutation assessor, GERP + +, etc.

## Result

### DNA concentration and PCR results

Qiaamp DNA FFPE tissue method (sample 1) and magnetic bead method (sample 2) were used to extract DNA samples for detection. OD260 / 280 of DNA samples extracted by QIAamp-DNA-FFPE tissue method is higher than 1.40, and the concentration is between 30.7ug/ul-44.5ug/ul. The extraction effect is relatively stable and the quality is relatively good. Od260 / 280 of the samples extracted by magnetic bead method was 1.3-3.76, the concentration was 12-22 UG / UL. The extraction effect fluctuated greatly.(Table 2)

**Table 2.** Concentration and OD value

|  | sample 1 |  | sample 2 |  |
| --- | --- | --- | --- | --- |
| No. | 1-1 | 1-2 | 2-1 | 2-2 |
| concentration (ug/ul) | 30.7 | 40.8 | 22 | 12 |
| OD | 1.48 | 1.44 | 1.31 | 3.76 |

The results of the first generation sequencing after gel cutting and purification of the bands (products of primer 1, primer 2, primer 4, primer 5, primer 6 and primer 7) that may meet the expected size were selected. The bands of two samples of primer 3 did not meet the expected size, and the two samples of primer 8 did not send the products for sequencing. We found that the PCR products of primer 1, primer 2 and primer 7 were expected to be sequenced successfully, while the PCR products of other primers, primer 5 and primer 6 were not sequenced successfully (no signal or very weak signal).(Fig.1)

The PCR products of primer 1, primer 2 and primer 7 were sequenced and analyzed by chromsa. Blast was compared with the reference genome sequence. The results showed that the positive sequence of primer 1 was 153bp. The reverse sequencing results showed that there were 107 BP matched reference sequences, and the rest of them had binding problems. The two-way sequencing results of primer 2 have all matched reference sequences. The two-way sequencing results of primer 7 of Mawangdui No.1 and No. In a word, eight pairs of primers have been designed, about two pairs of primers can get the available sequencing results, both of which are on mitochondrial DNA.(Fig.2)

### Quality control of library and sequencing

Two methods, magnetic bead method (labeled A1) and qiaamp DNA FFPE tissue method (labeled B1), were used to extract DNA from Mawangdui No. We use Agilent 2200 detector and d1000 reagent chip to test the quality control of the library, The main peak size of the library was in the range of 300-400 BP, and the mass concentrations were 22.1 ng / UL, 55 ng / UL, 7.18 ng / UL and 13.4 ng / UL, respectively. The final conclusion was pass, which showed that the library constructed was reliable in quality (Table 3 and Fig.3).

**Table 3.** Results of library quality inspection

| Sample name | No | 2200 detect results |  |  | conclusion |
| --- | --- | --- | --- | --- | --- |
|  |  | Peak bp | mass | Molar |  |
|  |  |  | concentration<br>ng/ul | concentration<br>nmol/l |  |
| Sample 1 | B1 | 368 | 55 | 230 | Pass |
| Sample 2 | A1 | 348 | 21.1 | 93.3 | Pass |

**Figure 3.**
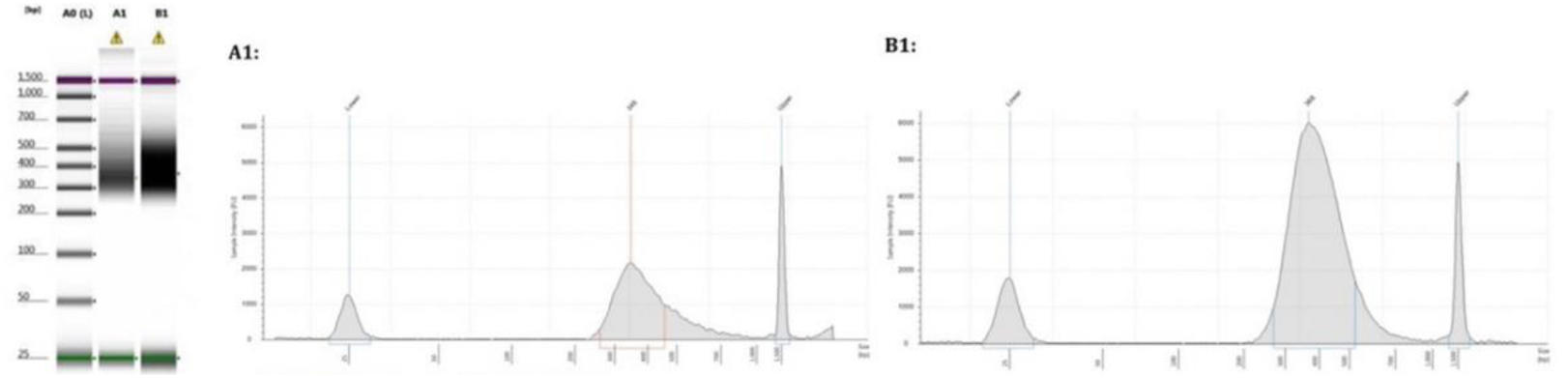
Strip chart and peak analysis chart of Agilent 2000 test results of two samples

The whole genome of the library constructed by two samples was sequenced by using illiumina 3000 sequencing platform, and about 90g of sequencing data (rawdata) were obtained. The quality control indexes Q20 and q30 showed that the sequencing data were effective. After filtering, the quality of 5’ and 3’ bases in reads is low, and the quality of bases in the middle part is high. The average quality of data after sequencing and filtering is high (Table 4). The quality of ancient DNA data will be evaluated by base content analysis, base quality analysis, base error rate analysis and damage plot. Base content is mainly used to detect whether at and GC are separated from sequencing data. In general, this phenomenon is introduced by database building or sequencing, and affects the subsequent analysis results of biological information. The base content distribution results show that the database building and sequencing show a good uniformity, which can be used for subsequent information analysis. From the distribution of base quality, the average quality of the data filtered by this sequencing is very high. The distribution of err rate of base shows that the overall error is at a low level. Damage plot shows the damage observed at the end of the reading and on the CpG island to confirm that the readings obtained are from ancient samples.

**Table 4.** Quality statistics of raw data

| Samples | Total reads | Total bases(bp) | GC% | Error rate(%) | Q20(%) | Q30(%) |
| --- | --- | --- | --- | --- | --- | --- |
| 1 | 10529890 | 1579483500 | 41.76 | 0.13 | 96.20 | 91.17 |
| 2 | 8147276 | 1222091400 | 41.99 | 0.42 | 95.18 | 89.56 |

Raw data is filtered according to the screening program of ancient DNA data analysis to generate high-quality sequence (clean data). After filtering, the quality of 5’ and 3’ bases in reads was low, and there was a little damage, which was consistent with the data quality characteristics of ancient DNA. The quality of the base in the middle part is high, and the average quality of the data filtered by this sequencing is high (Table 5).

**Table 5.** data filtering statistics

| Sample | HQ Reads | HQ Reads % | HQ Data (bp) | HQ Data % |
| --- | --- | --- | --- | --- |
| 1 | 10283624 | 97.66 | 1443785377 | 91.41 |
| 2 | 7957150 | 97.67 | 1093880284 | 89.51 |

### Data analysis

The testing sequence got about 1.5G data to analyze. The analysis of sequencing results needs to match the sequencing data to the reference genome. The reference genome sequence (hg19) data was downloaded from the UCSC database (http://genome.ucsc.edu/). The results of further sequence alignment showed that the total amount of data that could be compared to the reference genome accounted for 44.09-81.35% (Table 6). The whole gene coverage of 1x sequencing depth is about 0.10-0.35% (Table 7). Statistics of target sequencing regions, including total base of target region, total base of target region compared to reference genome target region, average sequencing depth and depth distribution, etc. (Fig. 4 and 5)

**Table 6.** Statistical results of sequence alignment

| Sample Name | Avg_read_len | ReadNum | 150rate (%) | Adapter (%) | reads mapped | bases mapped | reads_m<br>ap_rate | bases_m<br>ap_rate |
| --- | --- | --- | --- | --- | --- | --- | --- | --- |
| 1 | 150 | 8064898 | 76.59 | 23.41 | 7905943 | 1114464036 | 75.08 | 70.56 |
| 2 | 150 | 6389940 | 78.43 | 21.57 | 6627512 | 921194349 | 81.35 | 75.38 |

**Table 7.**
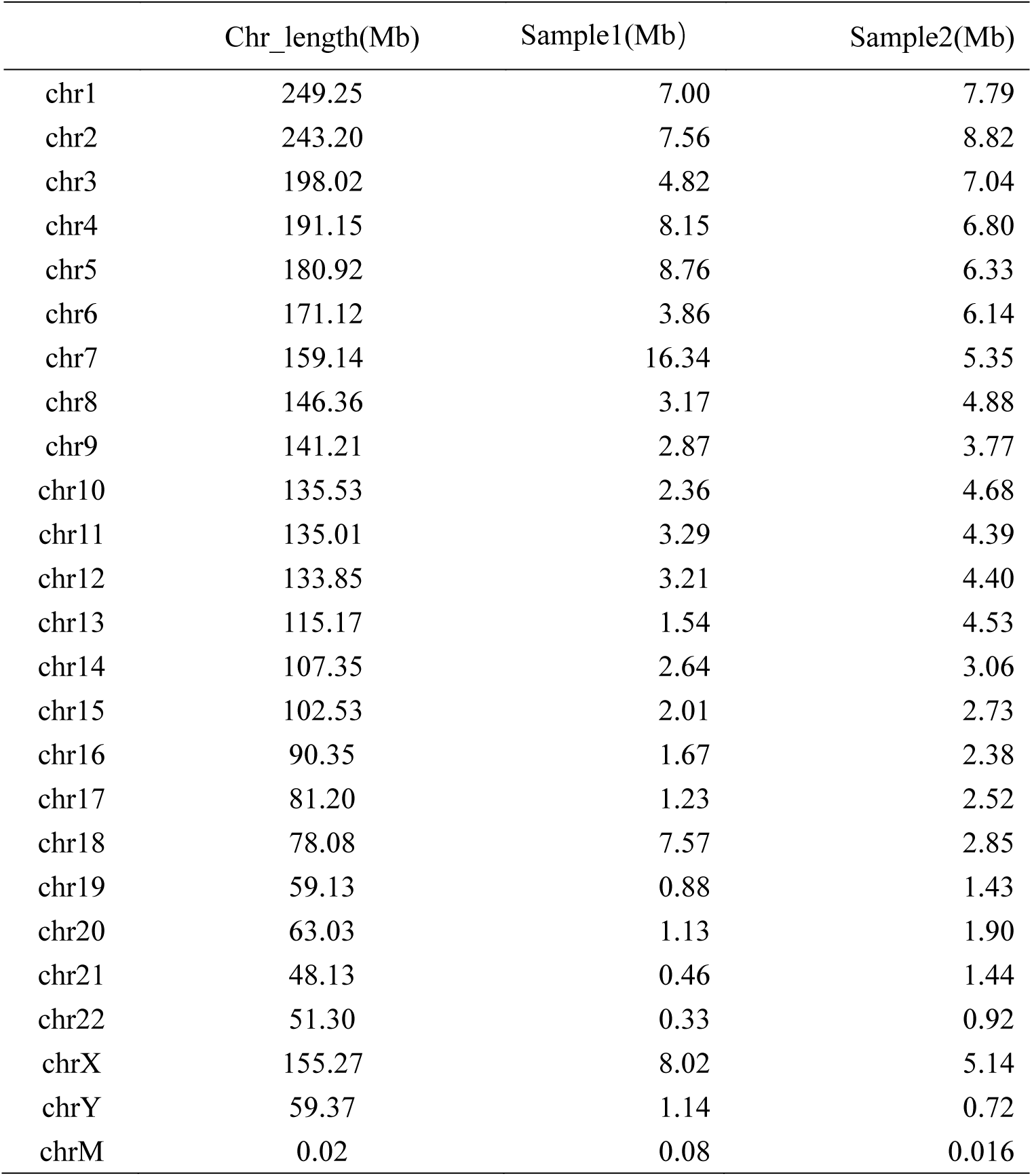
proportion results of chromosome comparison

**Figure 4.**
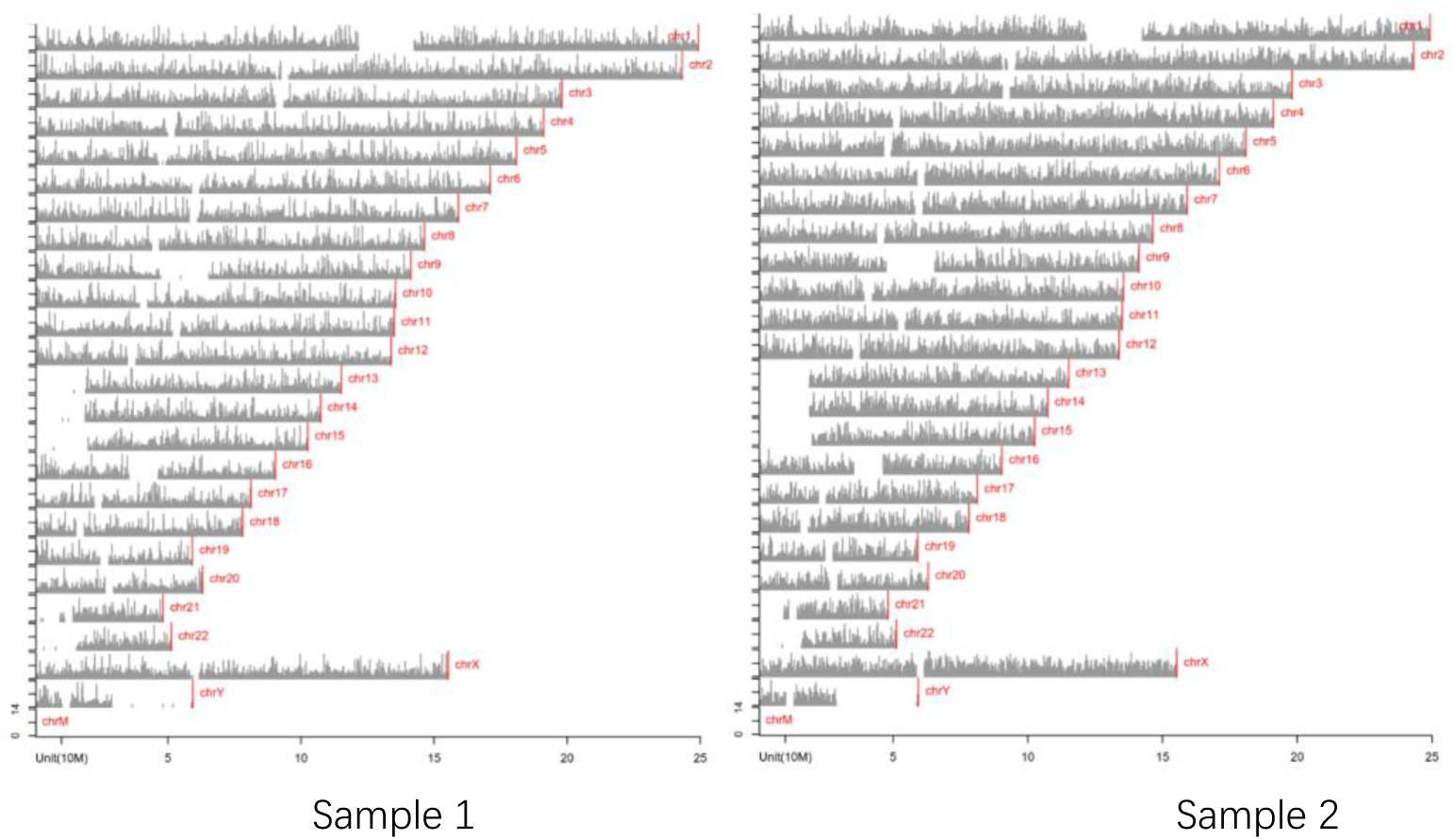
statistics of chromosome sequencing depth

**Figure 5.**
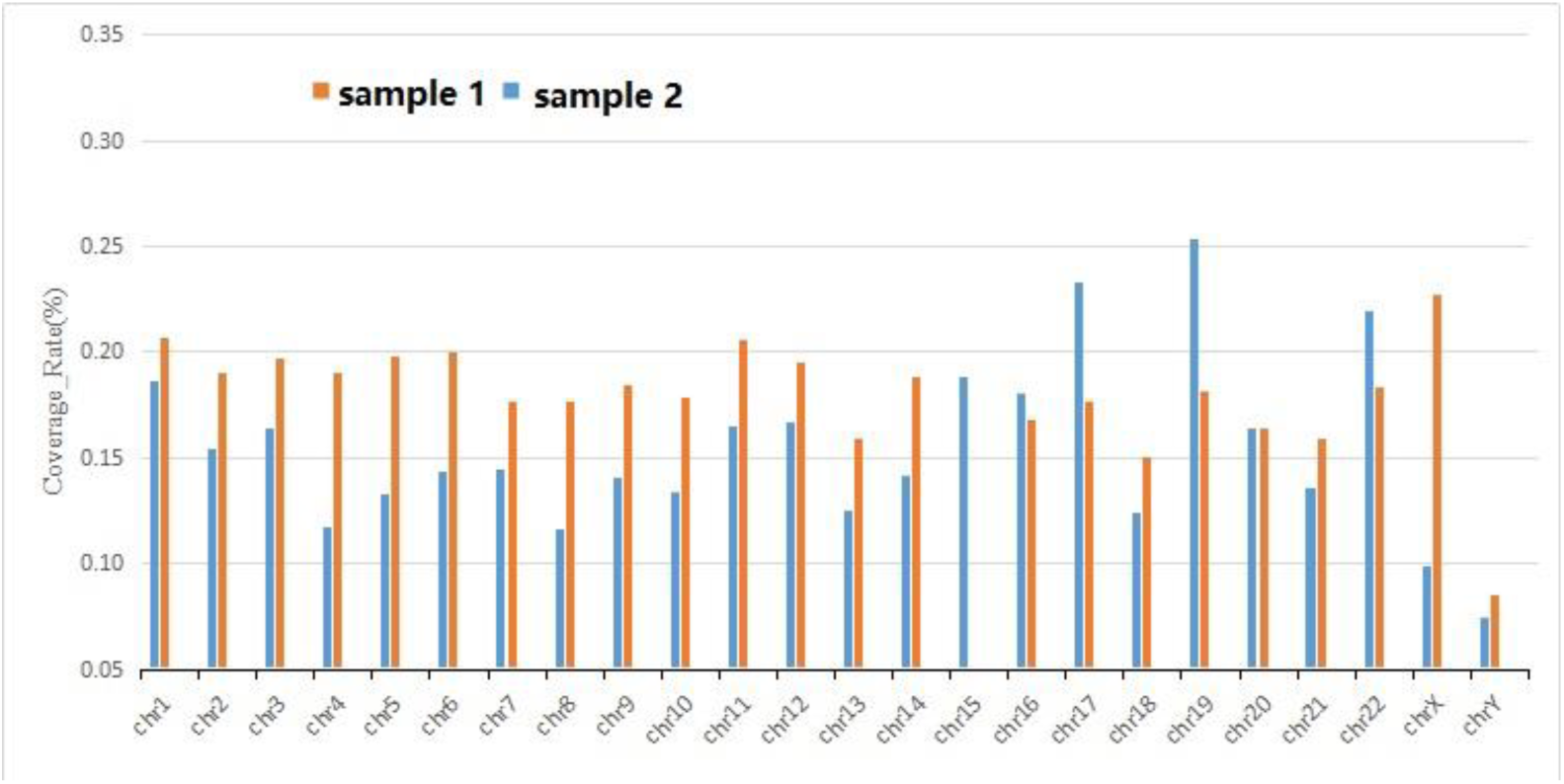
Chromosome coverage statistics

SNP (single nucleotide polymorphism) mainly refers to the DNA sequence polymorphism caused by the variation of a single nucleotide at the genomic level, including single base switching (TS), transversion (TV), etc. A total of 440 and 270 SNPs were covered by sample 1 and 2, respectively. The ratio of base conversion to base transversion (TS / TV) is 1.29and 1.78. (Table 8)

**Table 8.** Statistics of number and type of SNP

| Sample | Total sites | Heterozygous | Homozygous | Ts. | Tv. |
| --- | --- | --- | --- | --- | --- |
| Sample 1 | 440 | 89 | 351 | 247 | 193 |
| Sample 2 | 270 | 62 | 208 | 173 | 97 |

## Discussion

Generally, the samples in hot, humid and acid environment are not conducive to DNA preservation, and the subsequent formalin preservation solution is also harmful to DNA. However, Wang Guihai and Lu chuanzong, scientists of our country, have just unearthed the samples of Mawangdui female corpse. They have carried out the study of cell photometry and histochemical analysis of nucleic acid, which shows that the depolymerized and deteriorated nucleic acid components may be preserved in the nuclei of cartilage and liver of ancient corpse. It shows that it is possible to extract its DNA. The results also showed that although the DNA was not high in quality, concentration and purity, it might also contain a large number of proteins and other components, but PCR and first-generation sequencing all produced products. The results of the second generation high-throughput sequencing data show that there are still data available, although only about 1% of the whole genome.

Compare our SNP site information with the site detected in the wegene database (by Illumina Asian screening array Compared with 3 million SNP loci screened by beadchip (www.wegene.com), we found that only five SNP loci matched with the wegene database, which was far lower than the minimum requirement of ancestral analysis, rather than the analysis of population genetics, such as PCA, admixture, etc.

In conclusion, FFPE and high throughput sequencing methods is appropriate for DNA extraction of the ancient samples which were preserved in formalin, especially wet corpse.

## Acknowledgements

We thank all the members who contributed to this article. This paper was supported by frontier Research Program of Guangzhou Regenerative Medicine and Health Guangdong Laboratory. Grant No. 2018GZR110105020, The National Natural Science Foundation of China (31671211), and Science and Technology Planning Project of Guangdong Province, China (2017B030314056).

